# A novel meiotic protein required for homolog pairing and regulation of synapsis in *C. elegans*

**DOI:** 10.1101/2022.06.24.496392

**Authors:** Hyung Jun Kim, Abby F. Dernburg

## Abstract

Interactions between chromosomes and LINC (Linker of Nucleoskeleton and Cytoskeleton) complexes in the nuclear envelope (NE) promote homolog pairing and synapsis during meiosis. By tethering chromosomes to cytoskeletal motors, these connections lead to rapid, processive chromosome movements along the NE. This activity is usually mediated by telomeres, but in the nematode *Caenorhabditis elegans* special chromosome regions called “Pairing Centers” (PCs) have acquired this meiotic function. Through a genetic screen for mutations that cause meiotic nondisjunction, we discovered an uncharacterized meiosis-specific NE protein, MJL-1 (MAJIN-Like-1) that is essential for interactions between PCs and LINC complexes. MJL-1 colocalizes with PCs and LINC complexes during pairing and synapsis. Mutations in MJL-1 disrupt these interactions and eliminate active chromosome movements. *mjl-1* mutants display promiscuous nonhomologous synapsis, reduced clustering of PCs, and severely impaired homolog pairing. MJL-1 likely interacts directly with SUN-1 and DNA-binding proteins to connect PCs to the LINC complex. Similarities in the molecular architecture of chromosome-LINC complex attachments between *C. elegans* and other organisms suggest that these connections may play previously unrecognized roles during meiosis across eukaryotes.

## Introduction

Sexual reproduction relies on meiosis, the specialized cell division program that produces haploid gametes. During meiosis, homologous chromosomes must pair, synapse, and undergo crossover recombination to segregate accurately. Upon meiotic entry, each replicated chromosome is assembled into an array of loops anchored to a linear structure known as the chromosome axis (Zickler and Kleckner, 1999; Blat et al., 2002). Pairing of homologs is gradually stabilized by assembly of a protein matrix, the synaptonemal complex (SC), between axes (Zickler and Kleckner, 1999; Page and Hawley, 2004; Cahoon and Hawley, 2016). SCs promote and regulate crossover recombination, which results in chiasmata, physical linkages between homologous chromosomes that persist until segregation and mediate bipolar alignment on the spindle (Zickler and Kleckner, 2015; Page and Hawley, 2004; Kleckner 2006).

Chromosome pairing, synapsis, and recombination are promoted by nuclear envelope (NE)-associated chromosome dynamics during meiosis (Conrad et al., 1997; Chua et al., 1997; Cooper et al., 1998; Ding et al., 2004; Ding et al., 2007; Conrad et al., 2008). At meiotic entry, chromosomes become tethered to LINC (Linker of Nucleoskeleton and Cytoskeleton) complexes comprised of SUN and KASH domain proteins that span the two membranes of the NE (Hiraoka and Dernburg, 2009; Link and Jantsch, 2019). Cytoskeletal motors interact with LINC complexes on the cytoplasmic face of the NE, resulting in dramatic chromosome movements during early meiotic prophase (Hiraoka, 1998; Chikashige et al., 1994; Scherthan et al., 1996). This often leads to clustering of chromosome ends near cytoplasmic microtubule organizing centers to form a chromosome configuration known as the “meiotic bouquet” (Zickler and Kleckner, 1998; Scherthan, 2001; Hiraoka, 1998).

In the nematode *Caenorhabditis elegans*, specialized regions on each chromosome known as “Pairing Centers” (PCs) mediate homolog pairing and synapsis (Rog and Dernburg, 2013; Rosenbluth and Baillie 1981; McKim et al., 1988; Villeneuve 1994; MacQueen et al., 2005). Each PC recruits one of four meiosis-specific zinc finger proteins, ZIM-1, ZIM-2, ZIM-3, or HIM-8, through DNA binding sites present in clusters throughout the PC regions (Phillips et al., 2005; Phillips and Dernburg 2006, Phillips et al., 2009). During early meiotic prophase, PCs associate with LINC complexes comprised of SUN-1 and ZYG-12 (Penkner et al., 2007; Sato et al., 2009). ZYG-12 interacts with cytoplasmic dynein and perhaps other microtubule motors to drive processive movement of chromosomes that promote pairing and synapsis (Sato et al., 2009; Penkner et al., 2009; Wynne et al., 2012; Baudrimont et al., 2010).

In addition to the LINC complexes, additional meiosis-specific inner NE proteins are required for telomere-led chromosome movements in fission yeast and mice (Chikashige et al., 2009; Shibuya et al., 2015). Homologs of the mouse protein MAJIN (Membrane-Anchored Junction Protein) have been identified by sequence homology in many metazoans, but not in nematodes (da Cruz et al., 2020). Here we report the identification and characterization of a novel meiosis-specific NE protein that is essential for homolog pairing and synapsis in C. elegans. Based on the functions we have characterized, we named it MJL-1 (MAJIN-Like-1).

## Results

### Identification of MJL-1, a meiosis-specific NE protein

In *C. elegans*, defects in meiosis result in a High incidence of males (Him) phenotype due to nondisjunction of the *X* chromosome (Hodgkin et al., 1979); most meiotic mutants also produce many inviable embryos due to autosomal aneuploidy. We used a “Green eggs and Him” screen, based on a xol-1p::gfp reporter expressed in *XO* (male) embryos, to identify hermaphrodites with elevated meiotic nondisjunction (Kelly et al., 2000). Molecular lesions in the mutants were identified by outcrossing to the CB4856 Hawaiian strain, reisolating homozygous mutants, whole-genome sequencing of their progeny, and computational analysis to identify likely causal mutations (Doitsidou et al., 2010; Wicks et al., 2001; Swan et al., 2002). Most of the mutations we identified (50/52) were in genes previously shown to be important for meiosis, indicating that such Him screens are nearing saturation. Two mutations mapped to previously uncharacterized genes. One of these resulted in a premature stop codon in C17E4.4, which encodes a small protein with a single predicted transmembrane domain. Based on previous transcriptome data, C17E4.4 is specifically expressed in germline and arcade cells (Spencer et al., 2011; Han et al., 2017; Grun et al., 2014).

Using Cas9/CRISPR-based editing, we inserted an HA epitope tag at the N-terminus of the C17E4.4 protein. Hermaphrodites homozygous for this insertion produced a normal number of embryos and a slightly elevated frequency of male self-progeny (~1%, compared to 0.2% in wild-type broods). Immunofluorescence of the HA-tagged protein showed NE-specific localization throughout the meiotic region of the germline. The protein was undetectable in proliferating germline stem cells (GSCs) but was clearly observed at the NE upon meiotic entry and concentrated to form NE “patches” in transition zone (leptotene-zygotene) nuclei. Following synapsis, the protein redistributed throughout the NE and persisted until late pachytene (Fig. 1a, 1b).

**Figure 1.**
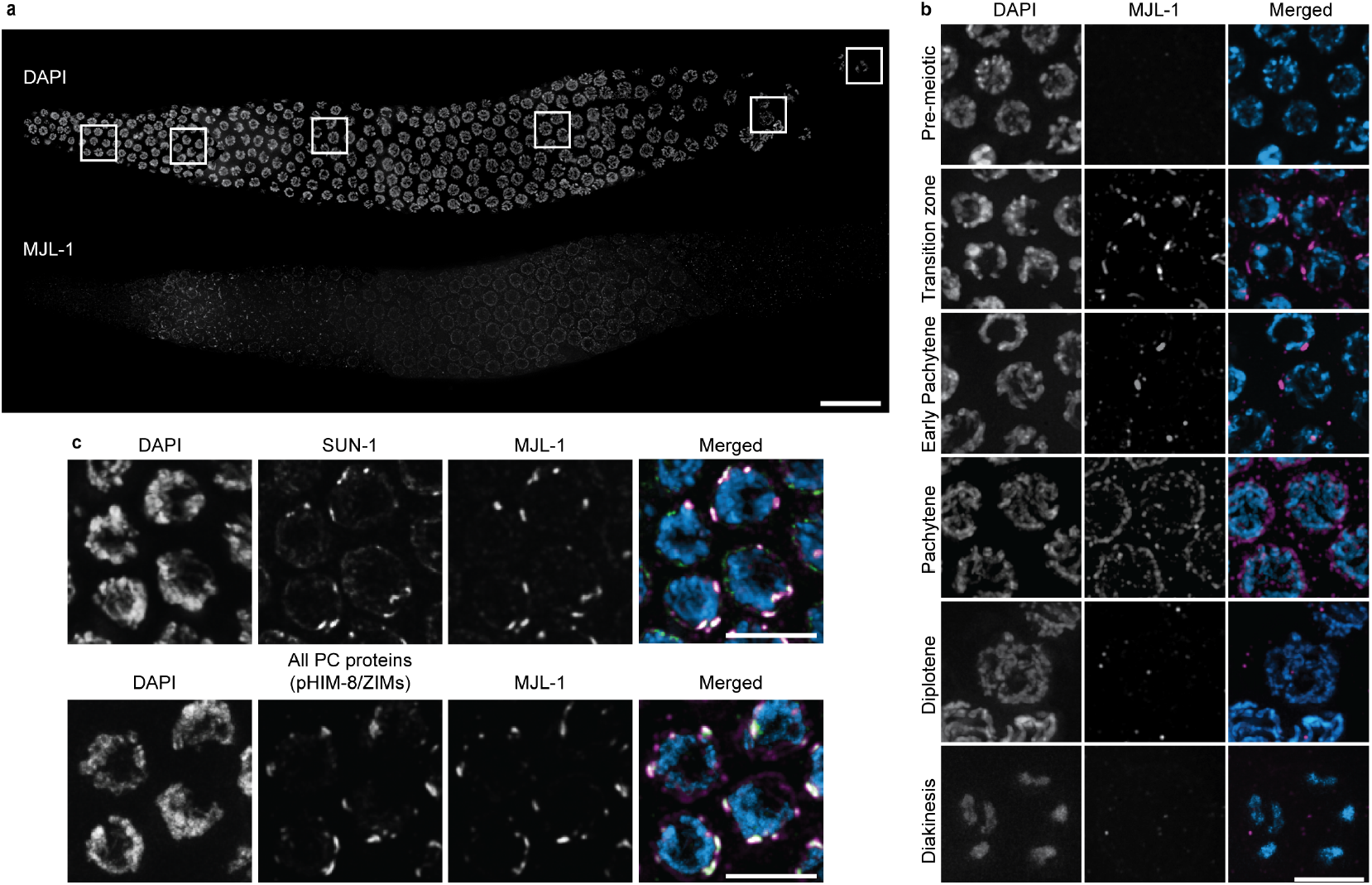
MJL-1 is a small NE protein that associates with PCs and LINC complexes. **(a)** Composite maximum-intensity projection images of whole gonads from *ha::mjl-1* hermaphrodites, stained with DAPI (top) and anti-HA antibodies (bottom). Scale bar, 20 μm. **(b)** Examples of nuclei at different stages of meiotic prophase. Scale bars, 5 μm. **(c)** MJL-1 colocalizes with phosphorylated PC proteins HIM-8 and ZIM-1, −2, and −3 (top) and SUN-1 (bottom) in *ha::mjl-1* hermaphrodites. PC proteins were detected using a phosphospecific antibody that recognizes these proteins when phosphorylated by CHK-2 (Kim et al., 2015). Scale bars, 5 μm.

During pairing and synapsis, the zinc finger proteins HIM-8, ZIM-1, −2, and −3 bind to PCs and interact with the LINC complex proteins SUN-1 and ZYG-12, which concentrate within the NE to form multiple patches. Immunofluorescence revealed that HA-tagged C17E4.4 colocalized with all four PC proteins during this transient stage (Fig. 1c).

Based on structural and functional similarities between C17E4.4 and mouse MAJIN (Shibuya et al., 2015), we named the gene *mjl-1*. Although MJL-1 shares no discernible sequence homology with MAJIN, both are small, single-pass transmembrane proteins with similar meiosis-specific functions (below) (Fig. 2a). MJL-1 is only weakly conserved within the genus *Caenorhabditis*, and we have not yet identified homologs in other nematode genera, including those that express homologs of the PC proteins (Fig. S1) (Rillo-Bohn et al., 2021).

**Figure 2.**
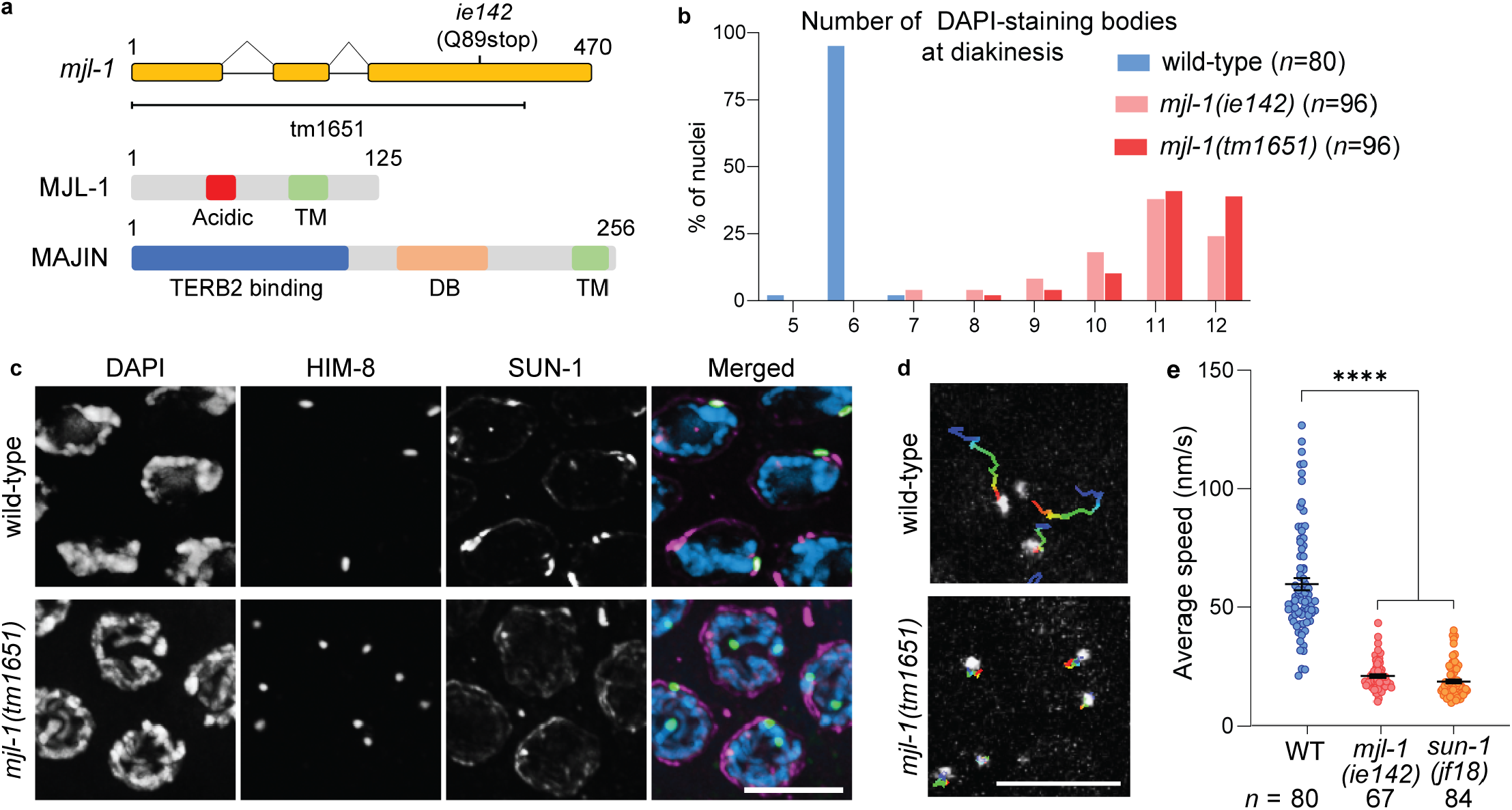
Loss of MJL-1 disrupts PC function. **(a)** Diagram of the *mjl-1* gene, indicating the mutations described in this work (top). Primary structure of MJL-1 in *C. elegans* and MAJIN in *M. musculus* (bottom) (TM: Transmembrane; DB: DNA binding domain). **(b)** Number of DAPI-staining bodies in oocyte nuclei at diakinesis in wild-type and *mjl-1* mutant hermaphrodites. **(c)** Loss of MJL-1 disrupts the connection between PC proteins and LINC complex. Transition zone nuclei were stained with antibodies against HIM-8 (green) and SUN-1 (magenta). Scale bars, 5 μm. **(d)** Projection of 75 s time course displacement track of GFP::HIM-8 in transition zone nuclei in wild-type and *mjl-1*(*ie142*). Scale bar, 5 μm. **(e)** Average speed of GFP::HIM-8 foci in transition zone nuclei in wild-type, *mjl-1*(ie142) (*p*<0.0001), and *sun-11*(jf18) (p< 0.0001) hermaphrodites. Each point represents a single nucleus. *p*-values were computed using Student’s *t*-test was used with Bonferroni *post-hoc* correction.

### MJL-1 is required for association of PCs with LINC complexes

The allele identified in our screen, *mjl-1*(*ie142*), is likely a null mutation since it contains a stop codon before its transmembrane domain. We also obtained a deletion allele, *mjl-1*(*tm1651*), from the Japanese National BioResource Project (NBRP) (Fig. 2a). Hermaphrodites homozygous for either mutant allele produced very few viable self-progeny (<1%), and among these, many were males (29% and 33%, statistically indistinguishable), indicative of extensive meiotic nondisjunction. At diakinesis, most oocytes in *mjl-1*(*ie142*) and *mjl-1*(*tm1651*) hermaphrodites displayed 10-12 DAPI-staining bodies (Fig. 2b), indicating that chromosomes failed to undergo crossing-over.

Based on its localization, we hypothesized that MJL-1 might mediate interaction between PCs and LINC complexes. Immunofluorescence confirmed that in *mjl-1* mutants, SUN-1 did not colocalize with PC proteins or form NE patches in transition zone nuclei (Fig. 2c). However, HIM-8 still appeared to associate with the NE, suggesting that HIM-8 may interact directly with the membrane or another NE protein (Fig. S2a). We crossed *mjl-1*(*ie142*) mutants to a strain expressing GFP-tagged HIM-8 (Wynne et al., 2012) to analyze chromosome movement. The average speed of HIM-8 foci in early meiotic nuclei was greatly reduced in the absence of MJL-1, from 59.8 nm/s in wild-type oocytes to 22.7 nm/s in *mjl-1*(*ie142*) (p<0.0001), similar to our measurements for *sun-1*(*jf18*) (18.7 nm/s) (Fig. 2d, 2e). Previous analysis has shown that *sun-1*(*jf18*), which results a missense mutation (G311V) in the SUN domain, abrogates active chromosome movement (Penkner et al., 2009; Baudrimont et al., 2010). The residual movement in *mjl-1*(*ie142*) and *sun-1*(*jf18*) mutants is likely due to diffusion rather than active motility (Wynne et al., 2012; Baudrimont et al., 2010; Woglar et al., 2013).

### MJL-1 is required to regulate synapsis

We observed extensive SC assembly despite very low levels of homolog pairing in *mjl-1* mutant nuclei (Fig. 3a, 3b). Similar promiscuous nonhomologous synapsis is observed in *sun-1*(*jf18*) mutants (Penkner et al., 2007; Sato et al., 2009). This effect of MJL-1 loss-of-function is consistent with prior evidence that the interaction between PCs and LINC complexes regulates synapsis so that it occurs only between homologous chromosomes. Nonhomologous synapsis in *mjl-1* and *sun-1*(*jf18*) mutants is more extensive than that caused by disruption of ZYG-12 or dynein (Sato et al., 2009; Zhang et al., 2015), suggesting a more direct role for these proteins in regulating synapsis initiation.

**Figure 3.**
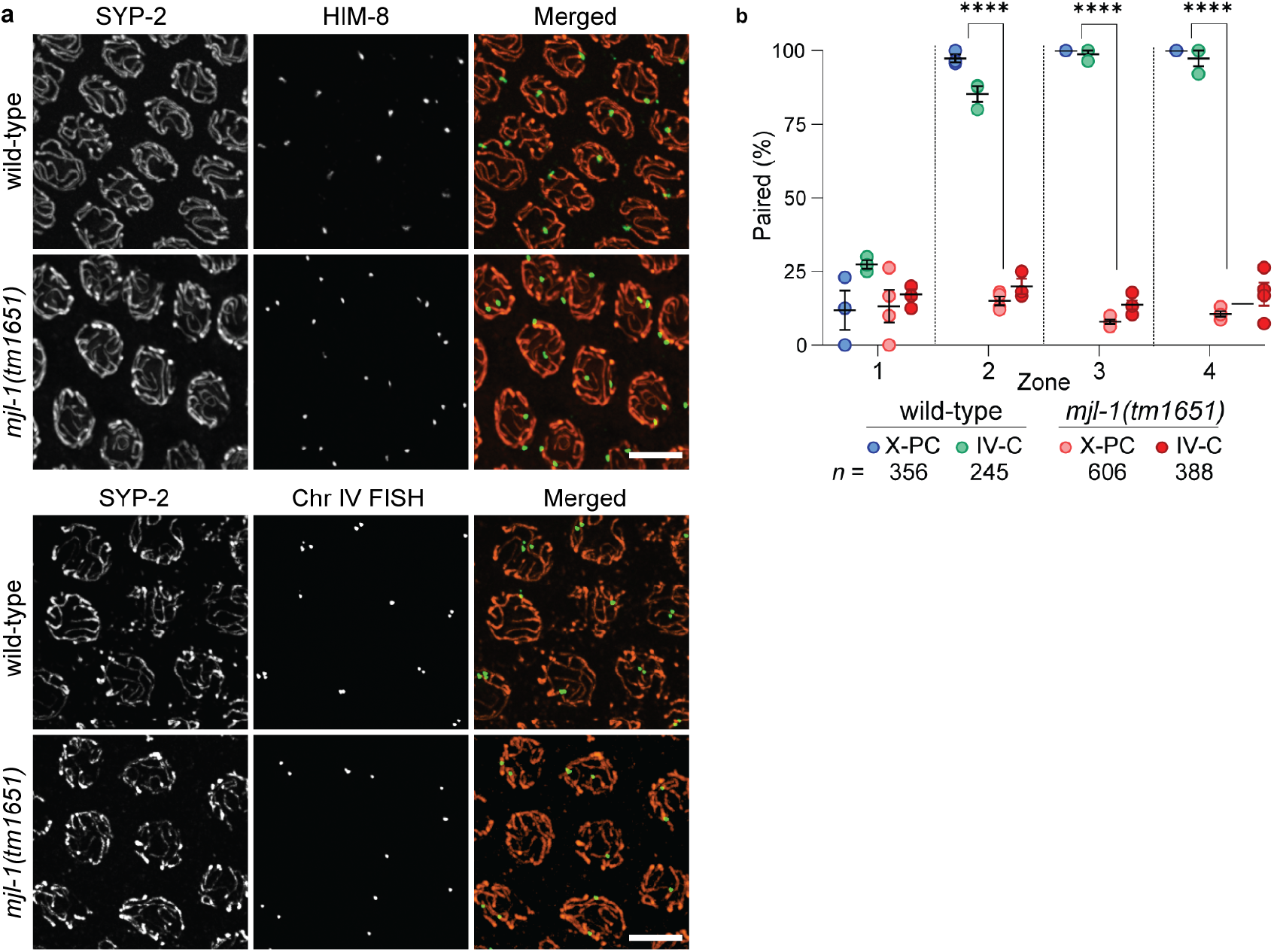
Deletion of *mjl-1* results in promiscuous nonhomologous synapsis. **(a)** Mid-pachytene nuclei stained with antibodies against HIM-8 (green) and SYP-2 (orange). Scale bars, 5μm. **(b)** Quantification of chromosome pairing in wildtype and *mjl-1*(*tm1651*) hermaphrodites using immunofluorescence (HIM-8) and FISH (Chr IV) *p*<0.0001). Gonads were divided into four zones (zone 1: pre-meiotic cells; zones 2-4: region spanning early prophase through pachytene, divided into three zones of equal length). p-values were calculated by one-way ANOVA with pairwise Bonferroni *post-hoc* correction.

### MJL-1 depends on SUN-1 for its NE localization

Unexpectedly, MJL-1 was not detected by immunofluorescence in *sun-1*(*ok1282*) null mutants or following auxin-induced degradation of SUN-1 (Fig. 4a). The abundance of MJL-1 detected on western blots was also strongly reduced following auxin-induced degradation of SUN-1 (Liu et al., 2021) (Fig. S2b). These findings indicate that SUN-1 is required for subcellular localization and/or stabilization of MJL-1. In contrast, neither the *sun-1*(*jf18*) missense mutation nor auxin-induced degradation of the KASH domain protein ZYG-12 (Liu et al., 2021) disrupted the NE localization of MJL-1 (Fig. 4a), suggesting that ZYG-12 is not required for association between MJL-1 and SUN-1.

**Figure 4.**
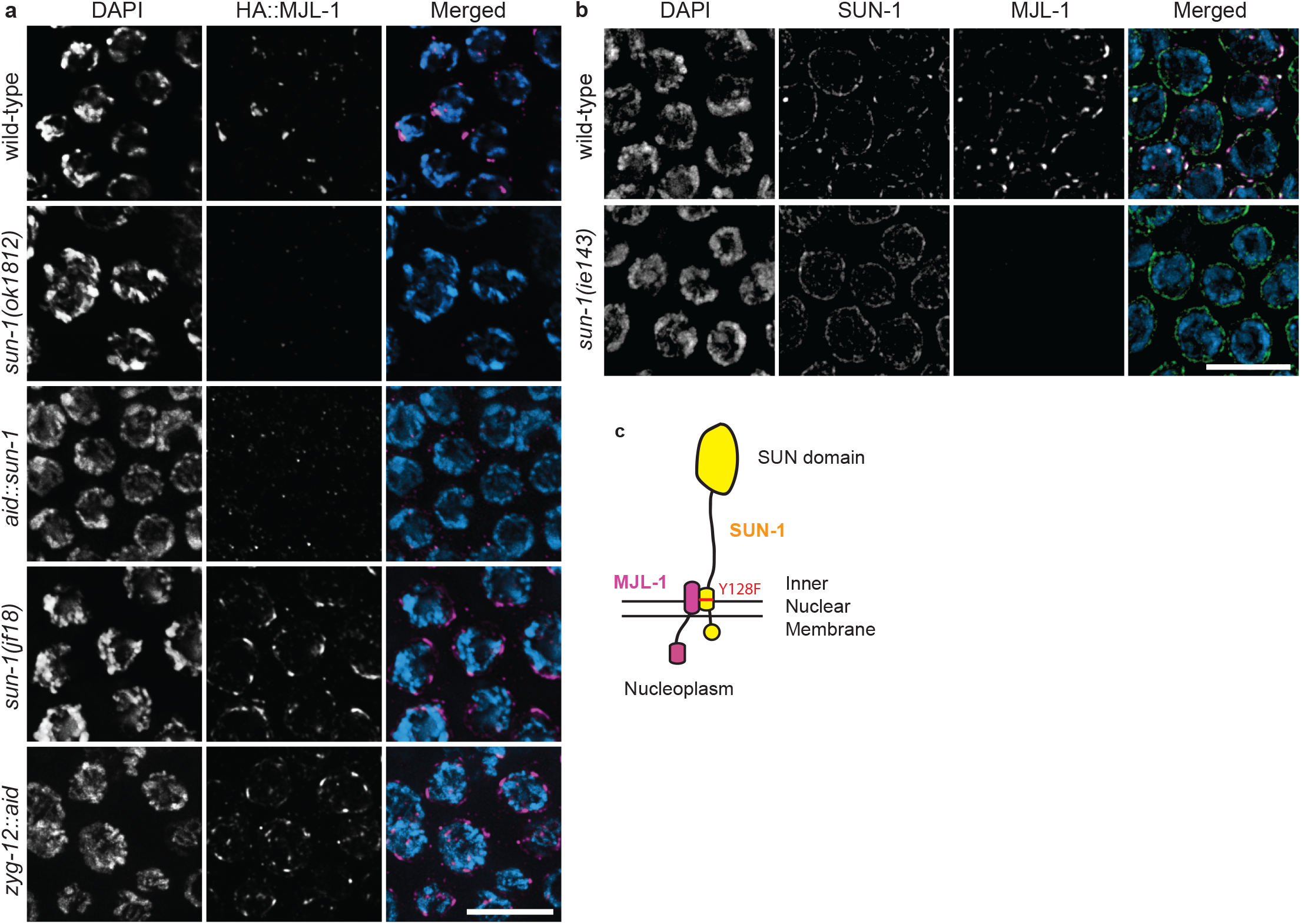
SUN-1 is required for NE localization of MJL-1. **(a)** Transition zone nuclei in wild-type and mutant hermaphrodites expressing HA::MJL-1, stained with anti-HA antibodies (magenta in merged images). Scale bar, 5 μm. **(b)** The Y128F mutation in *sun-1*(*ie143*) disrupts interaction between MJL-1 and SUN-1. Transition zone nuclei were stained with antibody against SUN-1 (green in merged images) and HA (magenta) from wild-type and mutant hermaphrodites expressing HA::MJL-1. Scale bar, 5 μm. **(c)** Illustration of the inferred interaction between MJL-1 and SUN-1.

Our screen also identified a novel meiosis-specific separation-of-function mutation in *sun-1* that resulted in phenotypes similar to *sun-1*(*jf18*). This new missense mutation, *sun-1*(*ie143*), changes tyrosine 128 within the predicted transmembrane domain of SUN-1, very close to the perinuclear domain, to phenylalanine (Y128F). In contrast to *sun-1*(*jf18*), *sun-1*(*ie143*) resulted in loss of MJL-1 protein in the germline, indicating that it disrupts the interaction between MJL-1 and SUN-1 (Fig. 4b, 4c), and thus suggesting that these proteins interact through their transmembrane and/or perinuclear domains. These regions in MJL-1 are conserved within *Caenorhabditis* (Fig. S3).

MJL-1 was also detected in apoptotic nuclei in the loop region of the germline. These nuclei retain SUN-1 at their NE but not ZYG-12 (Fig. S4), which may reflect disruption of the outer nuclear membrane during apoptosis, although this has not been directly demonstrated. The persistence of MJL-1 in these nuclei, together with our evidence that the protein requires SUN-1 for its localization to the NE (above) and may interact directly with PCs (below), indicates that MJL-1 probably resides within the inner nuclear membrane with its N-terminal domain in the nucleoplasm, similar to mouse MAJIN (Shibuya et al., 2015).

### Interactions between MJL-1 and PC proteins

Sequence alignment of *Caenorhabditis* MJL-1 homologs revealed a short region of relatively high conservation enriched for acidic residues (Fig. 5a). Using two crRNAs flanking this region, we generated an in-frame deletion of amino acids 34-49. The resulting MJL-1^Δacidic^ protein was expressed and localized to the NE but did not interact with PCs (Fig. 5a). Together with evidence that SUN-1 is essential for localization and stability of MJL-1 (above), this suggests that the mutant protein retains the ability to interact with SUN-1, but not with PC proteins. Homozygotes showed extensive nonhomologous synapsis, similar to *mjl-1* null mutants (Fig. S5). In contrast, an in-frame deletion of the sequence encoding amino acids 9-26, which are also relatively well conserved within *Caenorhabditis*, led to no apparent defects (data not shown).

**Figure 5.**
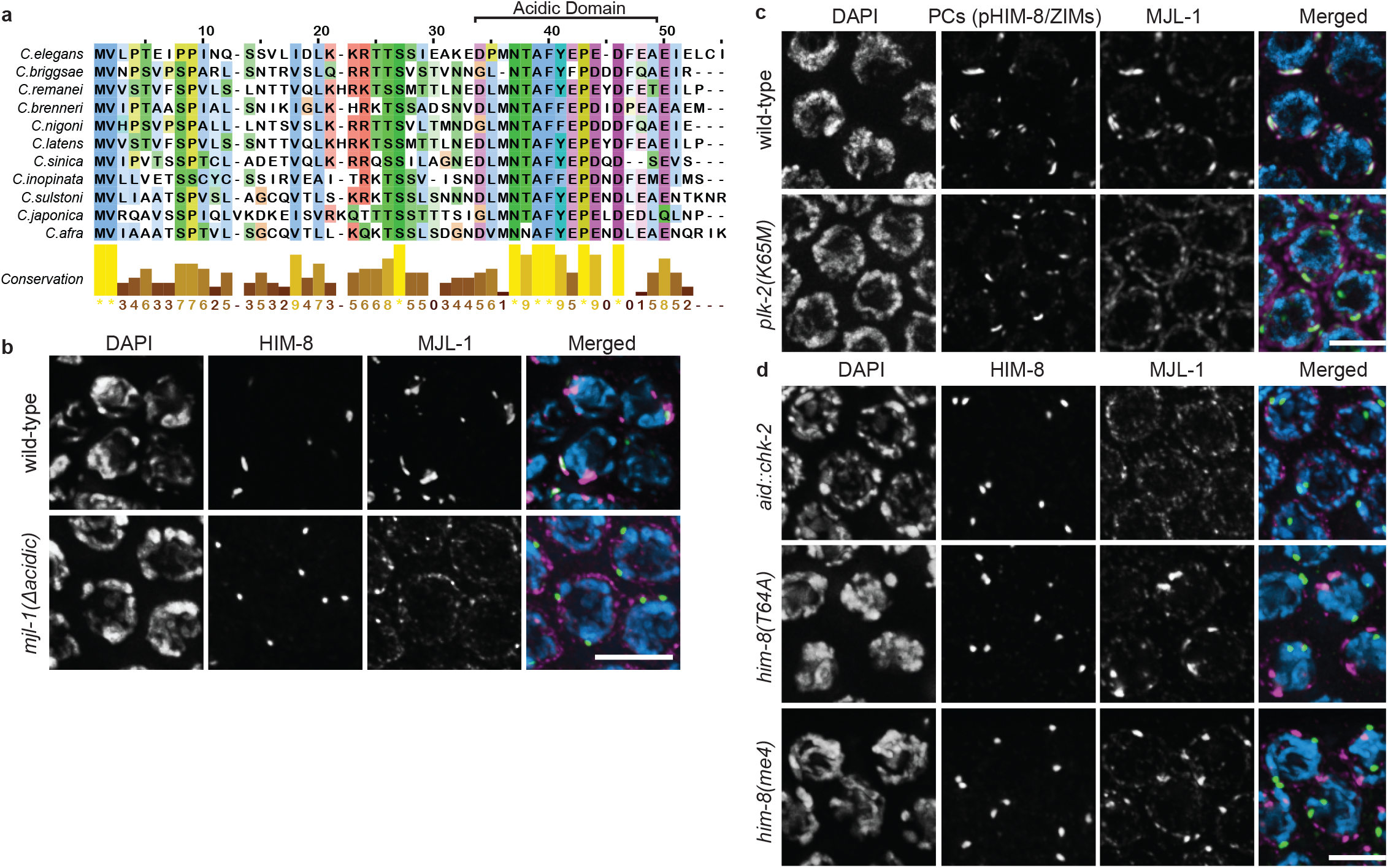
MJL-1 requires a small domain enriched in acidic amino acids to interact with PC proteins. **(a)** Sequence alignment of the N-terminal region of MJL-1 homologs within *Caenorhabditis*, generated using MAFFT. **(b)** Maximum intensity projection images showing zone nuclei stained with antibodies against HIM-8 (green in merged images) and HA (magenta) from hermaphrodites expressing HA::MJL-1WT (top) or HA::MJL-1^Δacidic^ (bottom). Scale bar, 5 μm. **(c)** PLK-2 activity is required for interaction between MJL-1 (magenta) and PC proteins (green). Scale bar, 5 μm. **(d)** Recruitment of PLK-2 by HIM-8 (green) is required for the association of HIM-8 with the MJL-1 (magenta). Scale bar, 5 μm.

The Polo-like kinase PLK-2 is recruited to PCs through Polo-box binding motifs in the zinc finger proteins and is required for interaction between PCs and the LINC complexes (Harper et al., 2011, Labella et al., 2011). Therefore, we asked whether PLK-2 activity at PCs is essential for the association of PC proteins with MJL-1. In a kinase-dead *plk-2*(K65M) mutant (Link et al., 2018; Brandt et al., 2020), MJL-1 localized throughout the NE and did not associate with PC proteins, suggesting that PLK-2 activity is required for the interaction between MJL-1 and PC proteins, but not for association between MJL-1 and SUN-1 (Fig. 5b), which is important for MJL-1 localization. Recruitment of PLK-2 to PCs requires phosphorylation of PC proteins by CHK-2 (Kim et al., 2015). We found that auxin-induced degradation of CHK-2 also abrogated the association between MJL-1 and PCs (Fig. 5c).

We examined the association of MJL-1 with two mutant versions of HIM-8, the PC protein specific for the X chromosome. HIM-8^T64A^, which does not recruit PLK-2 due to a point mutation in its Polo-box binding motif (Harper et al., 2011), and HIM-8^S85F^ (encoded by *him-8(me4*) allele), which fails to recruit both CHK-2 and PLK-2 (Kim et al., 2015), did not associate with patches of MJL-1 (Fig. 5c). Thus, PLK-2 activity at PCs is required for the interaction between PC proteins and MJL-1. We did not detect interactions between the putative nucleoplasmic domain of MJL-1 and PC proteins in a yeast two-hybrid assay (data not shown) but this may reflect a requirement for meiosis-specific post-translational modifications of these proteins.

### MJL-1 promotes homolog pairing

Inhibition of SC assembly prevents nonhomologous synapsis and partially restores X chromosome pairing in *sun-1*(*jf18*) mutants (Sato et al., 2009). To assess the role of MJL-1 in pairing, we compared the extent of X chromosome pairing in *zyg-12::aid* following auxin treatment, *sun-1*(*jf18*), and *mjl-1*(*ie142*), all in the absence of synapsis. Loss of MJL-1 resulted in more severe pairing defects than the absence of ZYG-12 or the *sun-1*(*jf18*) mutation, indicating that connection of chromosomes to MJL-1 and LINC complexes facilitates pairing even in the absence of rapid chromosome movements, although some “baseline” pairing (~25%) was still detected in mutants lacking MJL-1 (Fig. 6a, 6b). We also observed a reduction in nonhomologous associations between HIM-8 and other PCs in the absence of MJL-1 compared to *sun-1*(*jf18*) mutants, suggesting that MJL-1 promotes clustering of PCs (Fig. 6c, 6d), which may be required for efficient pairing of homologous PCs (Fig. 6e).

**Figure 6.**
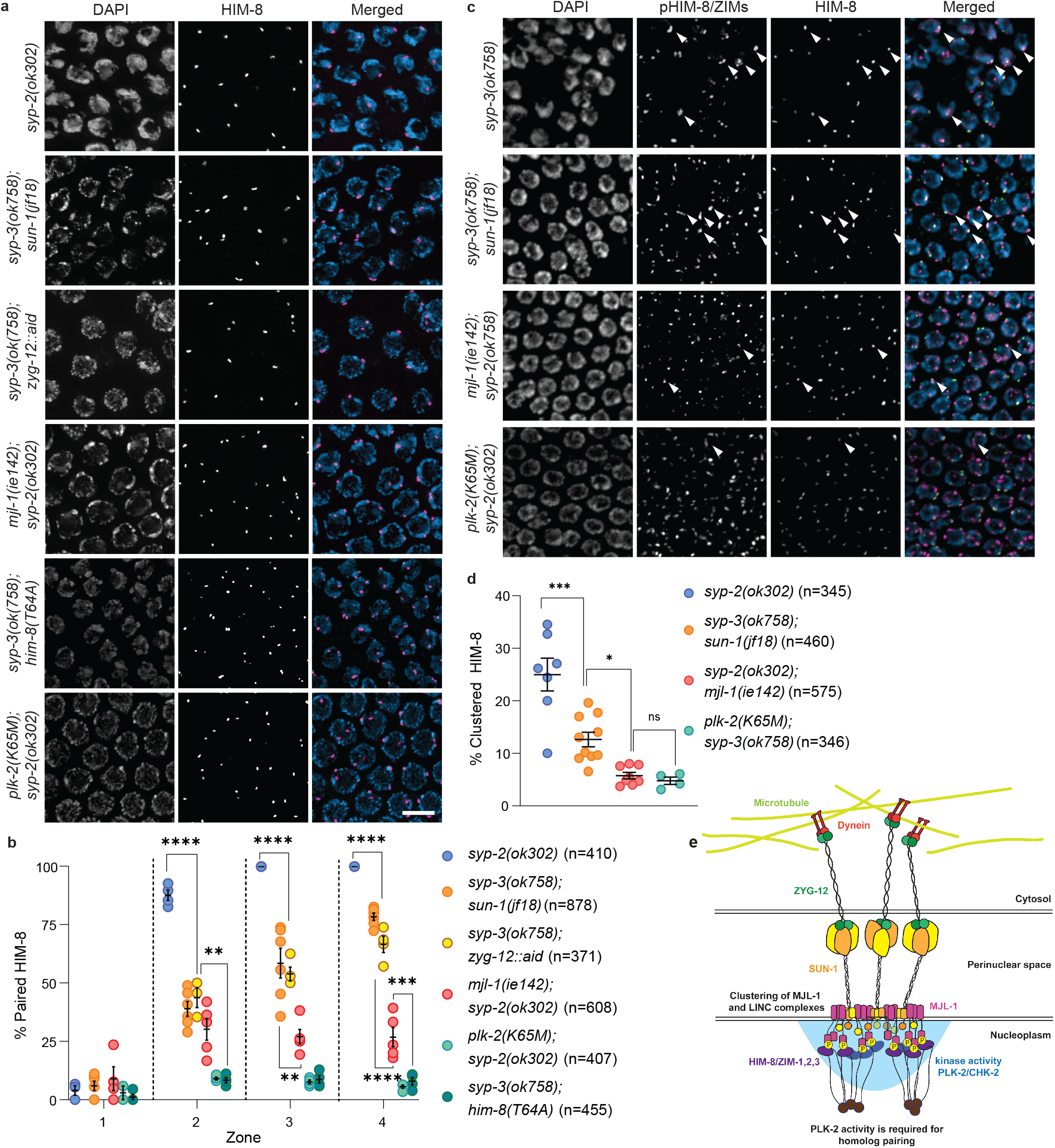
MJL-1 promotes pairing even in the absence of chromosome movements. **(a)** Blocking synapsis does not restore pairing of HIM-8 in *mjl-1*(*ie142*) mutants, in contrast to *sun-1*(*jf18*) or depletion of ZYG-12::AID. AID::SYP-3 was depleted by treatment with auxin for 24 h. Nuclei display polarized morphology due to lack of synapsis. Scale bar, 5 μm. **(b)** Quantification of X chromosome pairing. The extended transition zone was divided into three equal regions (zones 2-4) by length (zone 1: pre-meiotic). Each point represents a single gonad. *p*-values were calculated by one-way ANOVA with pairwise *post-hoc* Bonferroni correction (** p<0.01; *** p<0.001; **** p<0.0001). **(c)** In the absence of synapsis, clustering of HIM-8 with other PC proteins is lower in *mjl-1*(*ie142*) than in *sun-1*(*jf18*). Scale bar, 5 μm. **(d)** Quantification of clustering between HIM-8 and other PC proteins in various mutants. Only nuclei in zone 2 were analyzed, since pHIM-8/ZIM staining decreases in zone 3-4. *p*-values were calculated by one-way ANOVA with *post-hoc* pairwise Bonferroni correction (* *p* < 0.05; ** *p* < 0.01; *** p < 0.001; **** p < 0.0001)

### PLK-2 activity is required for homolog pairing

In contrast to *mjl-1* mutants, which showed a low level of X-chromosome pairing, hermaphrodites expressing kinase-dead PLK-2^K65M^ or HIM-8^T64A^ displayed almost no detectable *X* chromosome pairing in the absence of synapsis. This indicates that PLK-2 contributes to pairing beyond tethering PCs to MJL-1 during homolog pairing (Fig. 6a, 6b). Phosphorylation of the nuclear lamina by PLK-2 promotes chromosome mobility along the NE, likely by reducing homotypic interactions between LMN-1 proteins (Link et al., 2018). We found that depletion of LMN-1 did not rescue X chromosome pairing in *syp-3(ok758); him-8(T64A*) mutant, indicating that PLK-2 activity at PCs contributes to pairing through a lamin-independent mechanism (Fig. S6). In SC-deficient mutants, paired HIM-8 foci dissociated during late prophase, concomitant with the disappearance of PLK-2, indicating that PLK-2 activity increases affinity between homologous PCs (Fig. S7).

## Discussion

### Similarities and differences between MJL-1 and MAJIN

Homologs of MJL-1 are only detected within *Caenorhabditis*. PC proteins have been detected in related nematode genera (Rillo-Bohn et al., 2021), but show rapid divergence, particularly in their N-terminal domains, which act as scaffolds to recruit kinases and may also directly interact with NE proteins. If PC proteins interact with MJL-1, as suggested by our findings, rapid coevolution of these proteins may account for our inability to detect MJL-1 homologs in other nematode species.

Fission yeast Bqt3 and Bqt4 were the first NE proteins shown to be required for tethering of telomeres to LINC complexes during meiosis (Chikashige et al., 2009). MAJIN is likely to play a similar role in most metazoans (Shibuya et al., 2015; da Cruz et al., 2020). Bqt4, MAJIN, and MJL-1 share a similar structure, with a single transmembrane domain near their C termini, and may represent orthologous proteins despite their lack of sequence conservation (Hu et al., 2019; Shibuya et al., 2015).

Our evidence indicates that MJL-1 connects PC proteins to LINC complexes in *C. elegans*. Similarly, in mouse spermatocytes, MAJIN connects the meiosis-specific shelterin-binding proteins TERB1 and TERB2 to LINC complexes (Wang et al., 2020). MJL-1 requires PLK-2 activity to interact with PC proteins, while CDK2 activity is required for interaction between MAJIN and SUN1 in mice (Wang et al., 2020; Mikolcevic et al., 2016; Tu et al., 2017). In fission yeast, a direct interaction between NE proteins Bqt3 and −4 and LINC complexes has not been detected.

MAJIN and Bqt4 contain N-terminal DNA-binding motifs that are required for recruitment of telomeres to the NE (Hu et al., 2019; Shibuya et al., 2015). The DNA binding motif of MAJIN has been implicated in “telomere cap exchange,” whereby telomeres release shelterin and directly interact with TERB1, −2 and MAJIN (Shibuya et al., 2015). In contrast, MJL-1 lacks an apparent DNA-binding motif, and PCs in *C. elegans* associate with the NE even in the absence of MJL-1 or SUN-1.

### Roles of MJL-1 and LINC complexes in regulation of pairing and synapsis

Association of PCs with MJL-1 and LINC complexes inhibits inappropriate synapsis in *C. elegans* (Penkner et al., 2007; Sato et al., 2009). This is consistent with observations that these associations and resulting chromosome movements persist after pairing of homologs, which occurs soon after entry into meiosis. In contrast, loss of SUN-1 or MAJIN does not result in inappropriate synapsis in mice (Ding et al., 2007; Shibuya et al., 2015; Wang et al., 2019), which suggests that inhibition of nonhomologous synapsis by MJL-1 and LINC complexes may be unique to *C. elegans*. Nevertheless, CDK2, which inhibits inappropriate synapsis in mice, is also recruited to LINC complexes (Mikolcevic et al., 2016; Viera et al., 2009; Tu et al., 2017; Chen et al., 2021), suggesting that regulation of synapsis may be a general role of chromosome-LINC complex attachments, despite some differences in the details of this regulation between organisms.

Intriguingly, telomeric attachments in mammals and PC attachments in *C. elegans* each require a kinase (CDK2 and PLK-2, respectively) that is also involved in crossover regulation, suggesting that coordination between chromosome attachments sites and CO intermediates may be a conserved feature of meiosis (Woglar and Villeneuve, 2018; Zhang et al., 2021; Ashley and Rooij, 2001; Palmer et al., 2020).

## Acknowledgements

We thank Monica Colaiácovo and Verena Jantsch for providing antibodies, the Japanese National BioResource Project (NBRP) for the *mjl-1*(*tm1651*) mutant, and Yumi Kim for the *plk-2*(*K65M*) mutant. We thank members of the Dernburg lab for helpful comments on the manuscript and Chenshu Liu, Liangyu Zhang, and Fan Wu for strains. Some strains were provided by the CGC, which is funded by NIH Office of Research Infrastructure Programs (P40 OD010440). This work was supported by an Investigator award from HHMI to A.F.D. and a Fellowship from the Korea Foundation for Advanced Studies (KFAS) to H.J.K.

## Author contributions

Conceptualization: HJK, AFD

Methodology: HJK, AFD

Investigation: HJK

Supervision: AFD

Writing—original draft: HJK, AFD

## Materials and Methods

### *C. elegans* strains

N2 Bristol was used as the wild-type C. elegans strain; all mutations described here were generated in this background. The Hawaiian isolate CB4856 was used for genetic mapping. All strains were maintained at 20°C under standard laboratory conditions. The following mutations and balancers were used: *mjl-1(ie142), mjl-1(tm1651), sun-1(jf18), sun-1(ok1282), syp-2(ok307), syp-3(ok758), him-8(me4), him-8(T64A*) (Harper et al, 2011), *plk-2(K65M)::3xflag* (Brandt et al., 2020), *nT1[qIs51] (IV;V), hT2 [bli-4(e937) let-?(q782) qIs48] (I;III)*. The following constructs were used for auxin-inducible degradation: *zyg-12::ha::aid* and *aid::v5::sun-1* (Liu et al., 2021), *ha::aid::chk-2* (Zhang et al., 2021) where “aid” designates a 44-amino acid degron sequence (Zhang et al., 2015).

To generate *ha::mjl-1* strains, single-stranded (ss) DNA templates were designed to insert one or two copies of the HA tag at the N-terminus of MJL-1, separated by a flexible linker (GGGGS). These were co-injected with Cas9-NLS prebound to duplexed gRNAs, as well as a gRNA and ssDNA template for co-CRISPR of *dpy-10* (Arribere et al., 2014). (Final concentrations: *dpy-10* crRNA, 20 μM; *mjl-1* crRNA, 50 μM; trRNA, 40 μM; Cas9-NLS protein, 20 μM; *dpy-10* repair template, 1 μM, *mjl-1* repair template, 1 μM). To label MJL-1 using the split-GFP system and V5 tag, a template to insert GFP11 and V5 was co-injected with the Cas9-gRNA RNP complex into DUP223 *glh-1(sam129[glh-1::T2A::sGFP2(1-10)]*) (Goudeau et al., 2021). Essentially the same procedure was used to generate in-frame deletions in *mjl-1*, except that two gRNAs were used.

### Auxin-induced degradation

A stock solution containing 250 mM indole acetic acid (IAA, auxin) in EtOH was diluted to 1 mM in NGM agar just before pouring plates. After drying overnight, plates were seeded with *E. coli*. OP50 freshly cultured overnight to saturation in 50mL of LB was pelleted by centrifugation at 3,000xg for 5 min and resuspended in 500 μl of M9 buffer containing 1 mM auxin. This concentrated bacteria + auxin was spread on the plates and allowed to grow at room temperature for 1-2 days. To deplete degron-tagged proteins, young adult animals aged 24-48 h from L4 were picked onto these plates and analyzed 4-24 h later.

### RNA interference

Carbenicillin and IPTG were added to NGM agar to 200 μg/mL and 1 mM final just before pouring plates, respectively. Clones from the Ahringer laboratory (Fraser et al., 2000) were freshly cultured overnight to saturation in 10mL of LB containing 200 μg/mL carbenicillin. Then the culture was pelleted by centrifugation at 3,000xg for 5 min and resuspended in 50 μl of M9 buffer. Concentrated E. coli was spread on the plates and allowed to grow at 37°C for 1 day. For feeding, young adult animals aged 24-48 h from L4 were picked onto these plates and analyzed 24-48 h later. LMN-1 depletion was confirmed by shrunken nuclei in late pachytene-diplotene.

### Cytological Methods

Immunofluorescence and in situ hybridization were performed essentially as described previously (Dernburg et al., 1998). In brief: young adult worms were cut with a scalpel blade in egg buffer containing 0.05% tetramisole to release their gonads on slides, fixed in 1% formaldehyde in egg buffer for 2 min, transferred to tubes, and incubated with methanol pre-chilled to −30°C for 5 min. The tissue was then washed 3x in PBS containing 0.1% Tween 20 (PBST) at room temperature. Tissues were blocked using 1x Roche Blocking Reagent in PBST for 20 min. Primary antibodies were diluted into the same blocking solution and incubated with the tissues overnight at 4°C. Secondary antibodies were prepared in the blocking solution (1:200), mixed with samples, and incubated 1-2 h at room temperature. Samples were mounted in Prolong Diamond mounting medium containing DAPI (Invitrogen).

For fluorescence in situ hybridization, dissected gonads were fixed in 2% formaldehyde in egg buffer for 5 min, incubated with methanol pre-chilled to −30°C for 5 min, and washed 3x in 2x SSC containing 0.1% Tween 20 (2xSSCT) at room temperature. The tissue was then incubated in 50% formamide in 2x SSCT overnight at 37°C. 0.5-1 μl of 100μM fluorophore-conjugated oligonucleotide probes (IDT) “IV-2” (Adilardi and Dernburg, 2022) were added to 100 μl of hybridization buffer (50% formamide and 10% dextran sulfate in 2x SSCT) and tissues were moved into this mix, incubated 2-3 min at 91°C and then overnight at 37°C. The tissues were washed 3x in 2x SSCT at room temperature and mounted as for immunofluorescence (above).

Images were acquired using a DeltaVision Elite wide-field microscope system (GE) with a 100X 1.40 or 1.45 NA oil-immersion objective, or CSU-W1 SoRa confocal microscope system equipped with a 100X, 1.49 NA oil-immersion objective (Intelligent Imaging Innovations, Inc. [3i]). Deconvolution, projection, and analysis were performed using the softWoRx package or Slidebook 6 (3i).

### Antibodies

All antibodies used in this study were obtained from commercial sources or have been previously described. They include the following antibodies and dilutions: mouse monoclonal anti-HA (Invitrogen 2-2.2.14) (1:250), mouse monoclonal anti-V5 (Invitrogen 46-0705) (1:250), and rabbit polyclonal anti-V5 (Sigma V8137) (1:250). Custom polyclonal antibodies included rabbit anti-SUN-1 (1:250) (Sato et al., 2009), rat anti-HIM-8 (1:500) (Phillips et al., 2005), rabbit anti-SYP-2 (1:500) (Colaiácovo et al., 2003), rabbit anti-phospho-HIM-8/ZIMs (1:1000) (Kim et al., 2015). Fluorophore-conjugated secondary antibodies were purchased from Jackson ImmunoResearch and used at 1:200 dilution.

### Quantification of homolog pairing

3D distances between HIM-8 or FISH signals were measured using the “Measure Distance” tool in softWoRx. Foci closer than 0.6 μm were considered to be paired. We defined this threshold based on the maximum width of PC protein patches associated with paired chromosomes in wild-type oocytes. For analysis of pairing of FISH signals, we included only nuclei displaying two clear foci.

### *In vivo* imaging and quantification of chromosome movement

3D confocal image acquisition was performed essentially as described (Wynne et al., 2012) using Marianas spinning-disc confocal microscope system equipped with a 100X, 1.46 NA oil-immersion objective (Intelligent Imaging Innovations, Inc. [3i]). Exposure time was set to 100-150 ms depending on the brightness of foci. Stacks of 10–20 optical sections at 0.5 μm z-spacing were acquired every 5 s for a total of 5-10 min. 3D time-lapse images were analyzed using Imaris 9.2.0 (Bitplane). Background drift was corrected using the “Reference Frame” tool. Foci were detected using the “Spots” tool with an estimated XY diameter of 1.33 μm and filtered with “Quality” and “Intensity Sum”. Tracks were obtained with max distance of 1.75 μm and max gap of 3. Tracks from the background noise were manually removed. p-values were calculated using the pairwise t-test with post-hoc Bonferroni correction.

### Western blots

200 young adult animals aged 48 h from the L4 stage were picked into M9 buffer, washed three times, and then incubated in 1x SDS sample buffer at 50°C for 10 min. Samples were vortexed for 2–3 min until no visible solid material remained. Proteins were separated using SDS-PAGE gradient gels (Invitrogen NuPAGE™ 4-12%, Bis-Tris, 1.0 mm, 10-well) and transferred to Amersham Protran Nitrocellulose Membranes. The membrane was cut into slices and probed with mouse anti-HA (Invitrogen 2-2.2.14) (1:1,000) and mouse anti-α-tubulin (Sigma DM1A) (1:2,000) overnight at 4°C. HRP-conjugated donkey anti-mouse secondary antibody (Jackson ImmunoResearch) (1:10,000) was incubated with membranes 1-2 h at room temperature. Pierce ECL Western Blotting Substrate (Thermo) and SuperSignal West Femto Maximum Sensitivity Substrate (Thermo) were used as HRP substrates for detection of tubulin and 2xHA::MJL-1, respectively.

## Supplemental Figures

**Figure S1.**
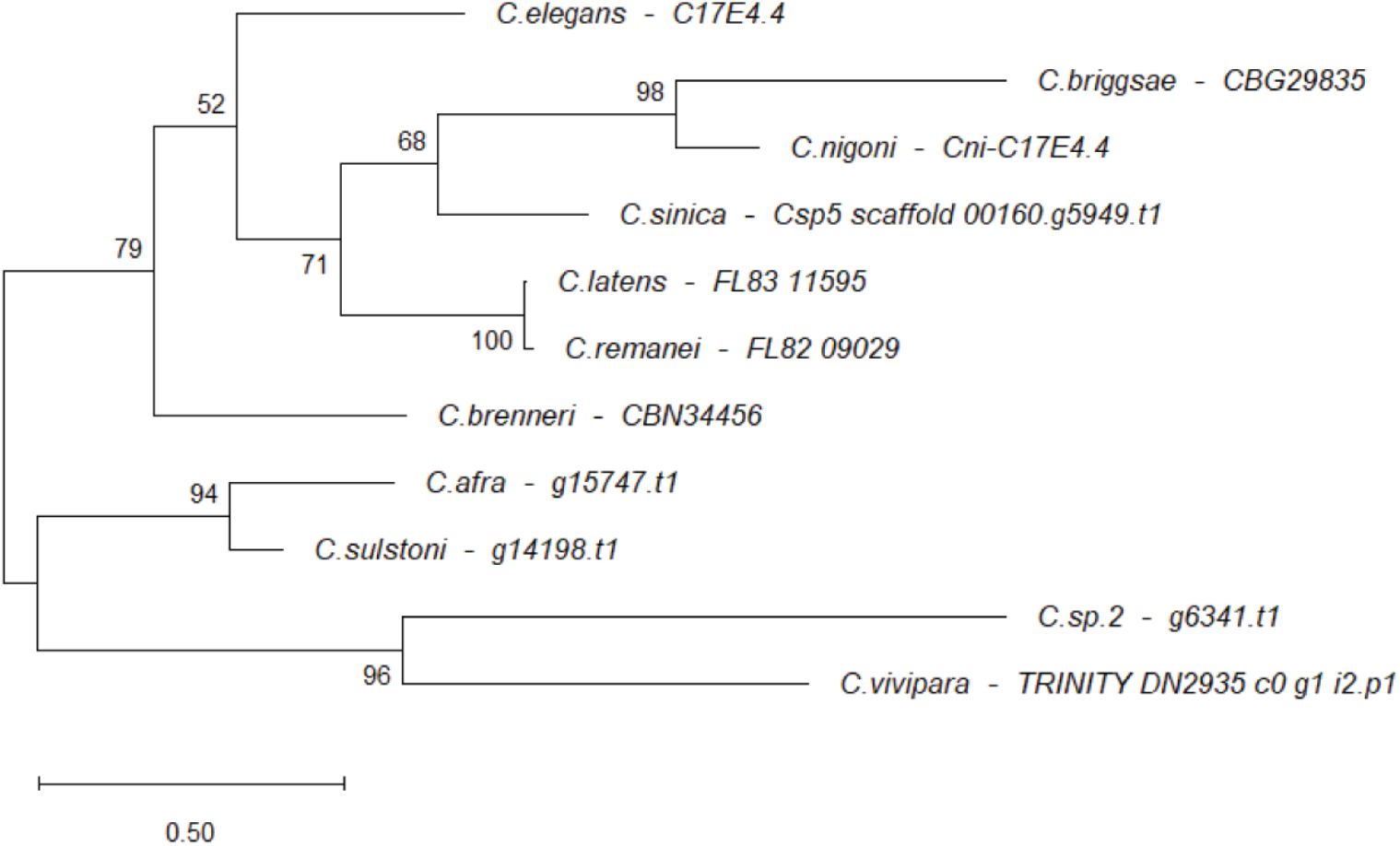
Phylogenetic tree of MJL-1 homologs in *Caenorhabditis*. A maximum-likelihood phylogenetic tree of MJL-1 homologs from representative *Caenorhabditis* species. Numbers on each node are Bootstrap values. Scale bar, 0.5 substitutions per site.

**Figure S2.**
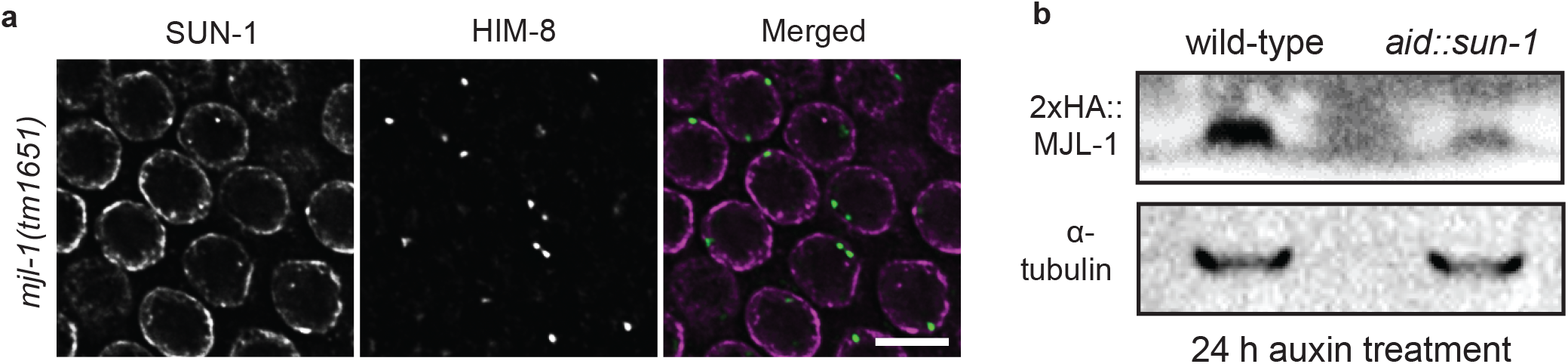
Interdependence among meiotic nuclear envelope proteins for localization and function. **(a)** Pairing Centers (PCs) interact with the nuclear envelope in the absence of MJL-1. A cross-section of meiotic nuclei showing immunofluorescence of SUN-1 (left), which marks the nuclear envelope in meiotic cells; HIM-8 (center) marks *X* chromosome PCs. Although pairing is severely reduced in *mjl-1*(*tm1615*), PCs are still associated with the nuclear envelope. **(b)** MJL-1 abundance is greatly reduced in the absence of SUN-1. Western blot of proteins in strains expressing 2xHA::MJL-1 and (degron-tagged) AID::SUN-1, showing the abundance of MJL-1 detected with anti-HA antibodies, either in the absence of auxin treatment or following depletion with auxin for 24 hours.

**Figure S3.**
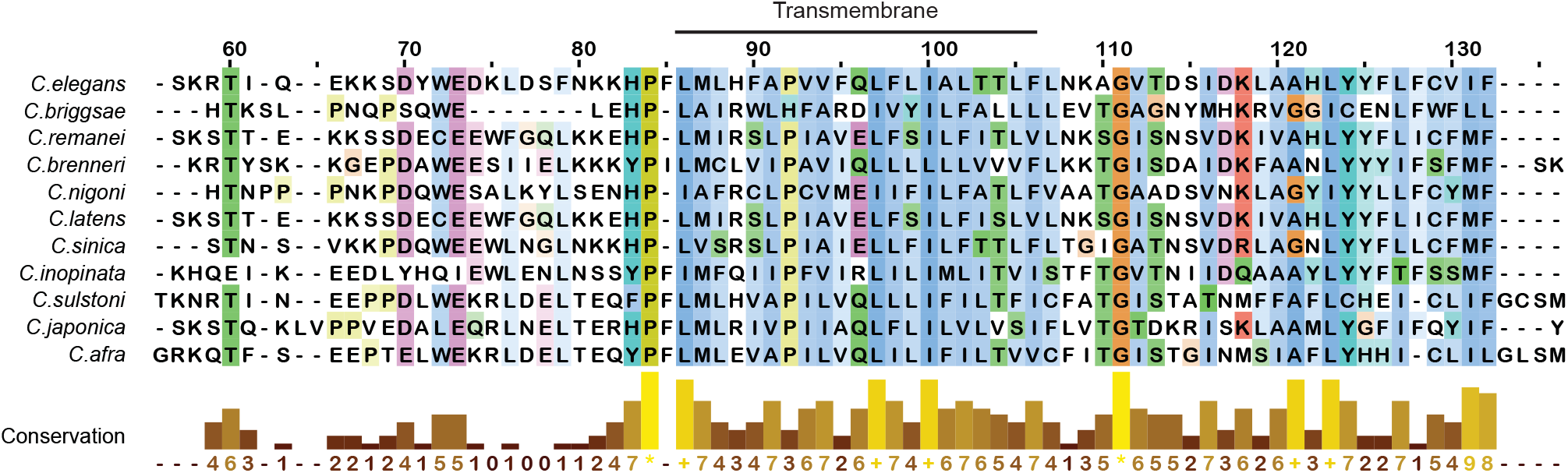
Sequence alignment of transmembrane and perinuclear regions in *Caenorhabditis* MJL-1 homologs. Alignment was generated using MAFFT.

**Figure S4.**
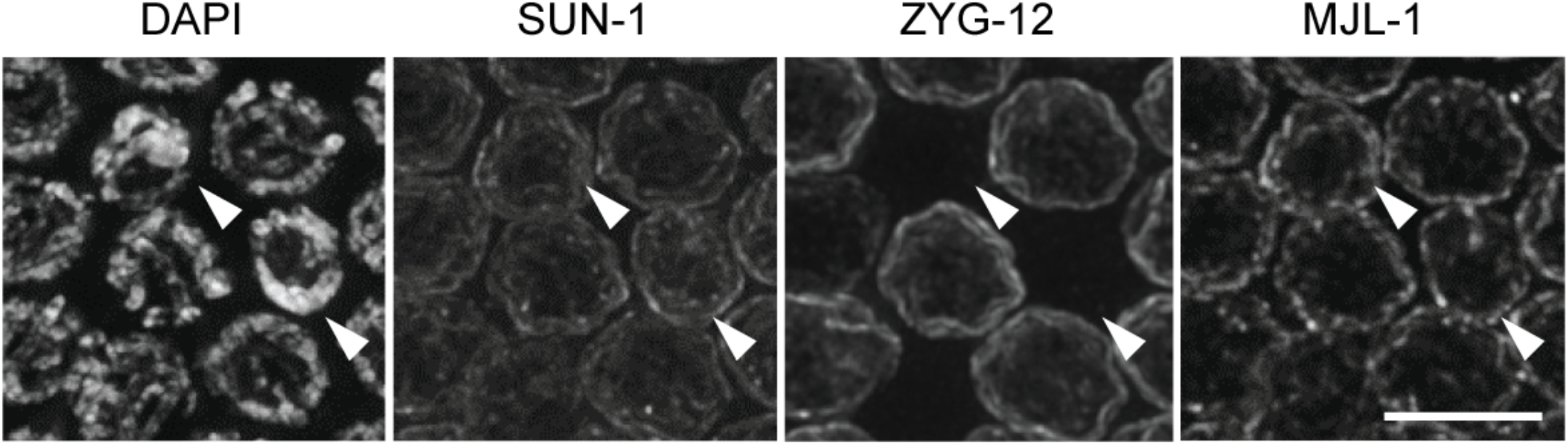
MJL-1 and SUN-1 are detected at the NE of oocyte nuclei undergoing apoptosis, while ZYG-12 is absent. Maximum-intensity projection images showing late pachytene nuclei. Arrowheads indicate apoptotic nuclei. Scale bar, 5 μm.

**Figure S5.**
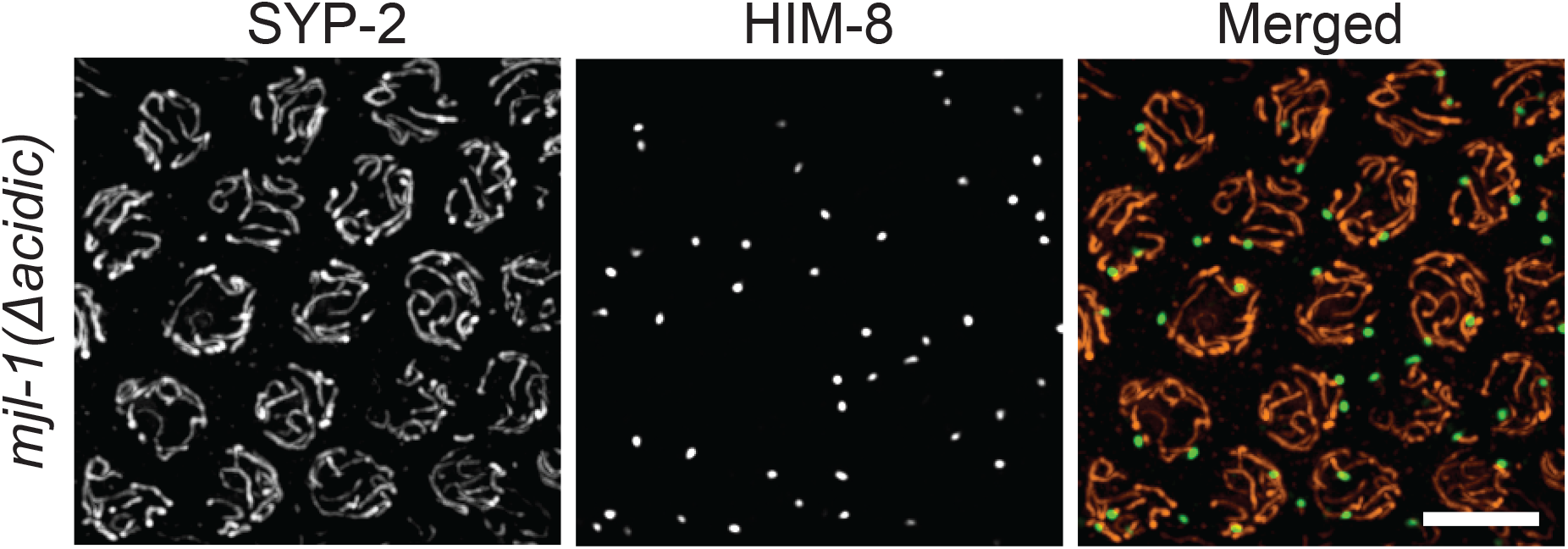
Deletion of a small acidic region in MJL-1 results in nonhomologous synapsis. Maximum-intensity projection images of pachytene nuclei stained with antibodies against HIM-8 (green in merged image) and SYP-2 (red). Scale Bar, 5 μm.

**Figure S6.**
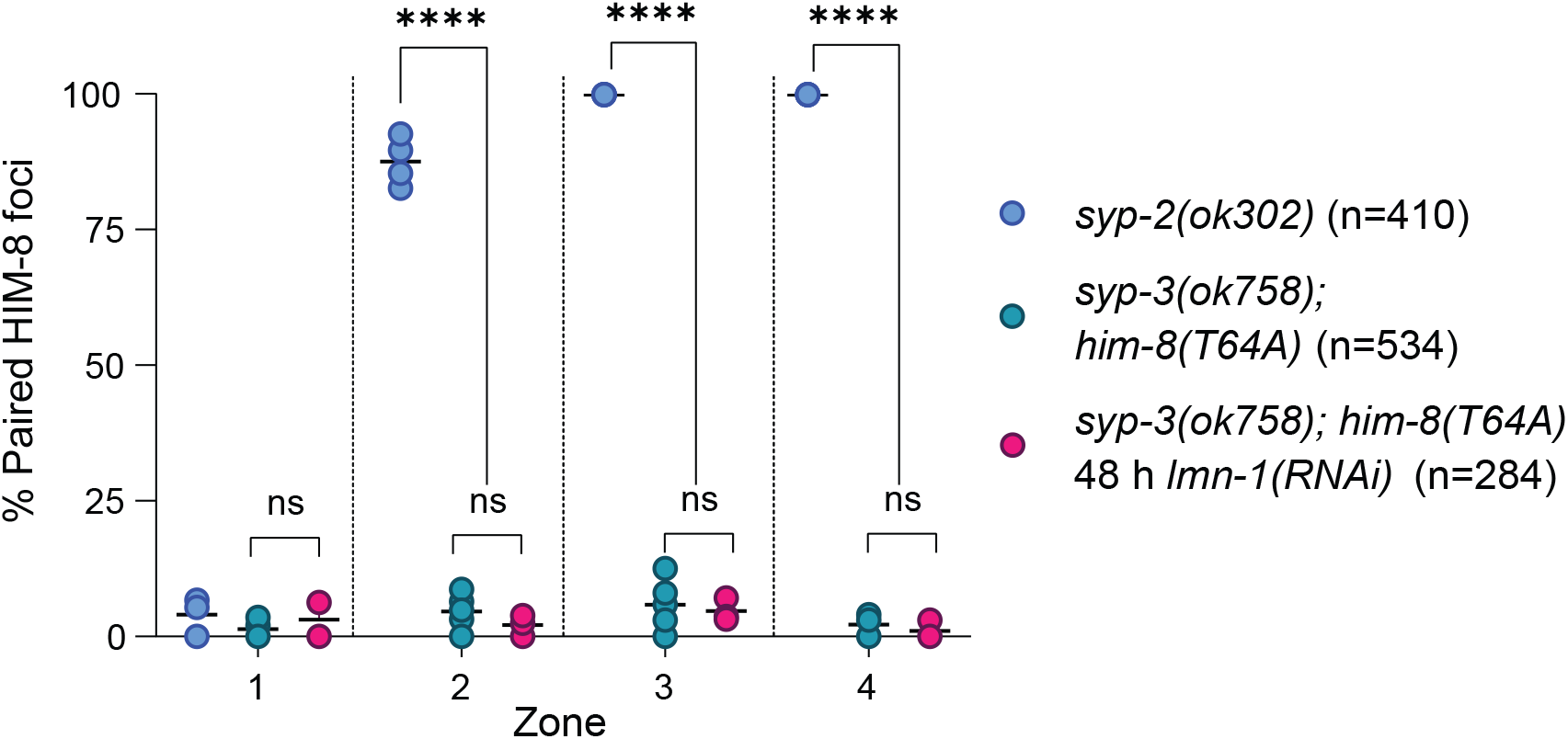
Depletion of LMN-1 by RNAi fails to rescue pairing in HIM-8^T64A^. Quantification of X chromosome pairing following depletion of the nuclear lamin protein LMN-1 by RNAi. HIM-8^T64A^ lacks a recruitment motif for PLK-2. The extended transition zone was divided into zones 2-4, with zone 1 corresponding to the pre-meiotic region. Each point represents a single gonad. *p-*values were calculated by one-way ANOVA with pairwise Bonferroni *post-hoc* correction (**** *p*<0.0001).

**Figure S7.**
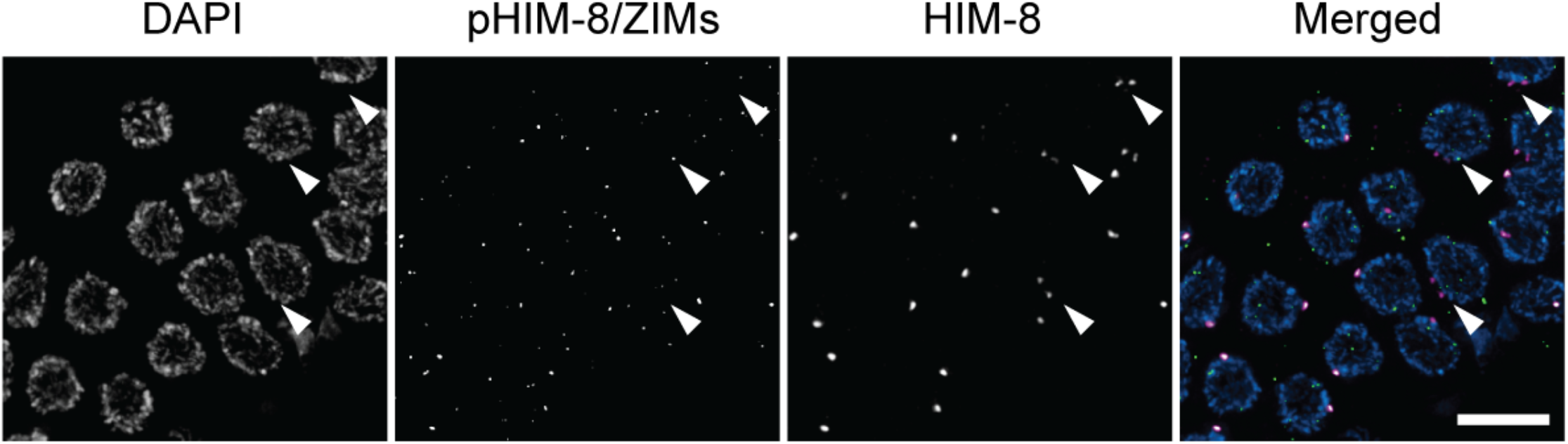
Loss of PLK-2 correlates with dissociation of synapsis-independent pairing. Images show maximum-intensity projections of the proximal region of the gonad, corresponding to the end of the extended transition zone, in *syp-2*(*ok307*) hermaphrodites. Gonads were stained with antibodies against phosphorylated PC proteins (green) and HIM-8 (magenta). Separation of HIM-8 foci correlates with a loss phosphorylation of HIM-8, indicative of loss of PLK-2 from the X chromosome PC.

## References

1. Adilardi, R. S. & Dernburg, A. F. Robust, versatile DNA FISH probes for chromosome-specific repeats in Caenorhabditis elegans and Pristionchus pacificus. G3 (Bethesda) (2022).

2. Arribere, J. A. et al. Efficient Marker-Free Recovery of Custom Genetic Modifications with CRISPR/Cas9 in Caenorhabditis elegans. Genetics 198, 837–846 (2014).

3. Ashley, T., Walpita, D. & de Rooij, D. G. Localization of two mammalian cyclin dependent kinases during mammalian meiosis. J Cell Sci 114, 685–693 (2001).

4. Baudrimont, A. et al. Leptotene/zygotene chromosome movement via the SUN/KASH protein bridge in Caenorhabditis elegans. PLoS Genet 6, e1001219 (2010).

5. Blat, Y., Protacio, R. U., Hunter, N. & Kleckner, N. Physical and functional interactions among basic chromosome organizational features govern early steps of meiotic chiasma formation. Cell 111, 791–802 (2002).

6. Brandt, J. N., Hussey, K. A. & Kim, Y. Spatial and temporal control of targeting Polo-like kinase during meiotic prophase. J Cell Biol 219 (2020).

7. Cahoon, C. K. & Hawley, R. S. Regulating the construction and demolition of the synaptonemal complex. Nat Struct Mol Biol 23, 369–377 (2016).

8. Chen, Y. et al. The SUN1-SPDYA interaction plays an essential role in meiosis prophase I. Nat Commun 12, 3176 (2021).

9. Chikashige, Y. et al. Telomere-led premeiotic chromosome movement in fission yeast. Science 264, 270–273 (1994).

10. Chikashige, Y. et al. Membrane proteins Bqt3 and-4 anchor telomeres to the nuclear envelope to ensure chromosomal bouquet formation. J Cell Biol 187, 413–427 (2009).

11. Chua, P. R. & Roeder, G. S. Tam1, a telomere-associated meiotic protein, functions in chromosome synapsis and crossover interference. Genes Dev 11, 1786–1800 (1997).

12. Colaiácovo, M. P. et al. Synaptonemal complex assembly in C. elegans is dispensable for loading strand-exchange proteins but critical for proper completion of recombination. Dev Cell 5, 463–474 (2003).

13. Conrad, M. N., Dominguez, A. M. & Dresser, M. E. Ndj1p, a meiotic telomere protein required for normal chromosome synapsis and segregation in yeast. Science 276, 1252–1255 (1997).

14. Conrad, M. N. et al. Rapid telomere movement in meiotic prophase is promoted by NDJ1, MPS3, and CSM4 and is modulated by recombination. Cell 133, 1175–1187 (2008).

15. Cooper, J. P., Watanabe, Y. & Nurse, P. Fission yeast Taz1 protein is required for meiotic telomere clustering and recombination. Nature 392, 828–831 (1998).

16. da Cruz, I., Brochier-Armanet, C. & Benavente, R. The TERB1-TERB2-MAJIN complex of mouse meiotic telomeres dates back to the common ancestor of metazoans. BMC Evol Biol 20, 55 (2020).

17. Dernburg, A. F. et al. Meiotic recombination in C. elegans initiates by a conserved mechanism and is dispensable for homologous chromosome synapsis. Cell 94, 387–398 (1998).

18. Ding, D., Yamamoto, A., Haraguchi, T. & Hiraoka, Y. Dynamics of Homologous Chromosome Pairing during Meiotic Prophase in Fission Yeast. Developmental Cell 6, 329–341 (2004).

19. Ding, X. et al. SUN1 Is Required for Telomere Attachment to Nuclear Envelope and Gametogenesis in Mice. Developmental Cell 12, 863–872 (2007).

20. Doitsidou, M., Poole, R. J., Sarin, S., Bigelow, H. & Hobert, O. C. elegans Mutant Identification with a One-Step Whole-Genome-Sequencing and SNP Mapping Strategy. PLoS ONE 5, e15435 (2010).

21. Fraser, A. G. et al. Functional genomic analysis of C. elegans chromosome I by systematic RNA interference. Nature 408, 325–330 (2000).

22. Goudeau, J. et al. Split-wrmScarlet and split-sfGFP: tools for faster, easier fluorescent labeling of endogenous proteins in Caenorhabditis elegans. Genetics 217, iyab014 (2021).

23. Grün, D. et al. Conservation of mRNA and Protein Expression during Development of C. elegans. Cell Reports 6, 565–577 (2014).

24. Han, S. et al. Mono-unsaturated fatty acids link H3K4me3 modifiers to C. elegans lifespan. Nature 544, 185–190 (2017).

25. Harper, N. C. et al. Pairing centers recruit a Polo-like kinase to orchestrate meiotic chromosome dynamics in C. elegans. Dev Cell 21, 934–947 (2011).

26. Hiraoka, Y. Meiotic telomeres: a matchmaker for homologous chromosomes. Genes Cells 3, 405–413 (1998).

27. Hiraoka, Y. & Dernburg, A. F. The SUN Rises on Meiotic Chromosome Dynamics. Developmental Cell 17, 598–605 (2009).

28. Hodgkin, J., Horvitz, H. R. & Brenner, S. Nondisjunction Mutants of the Nematode CAENORHABDITIS ELEGANS. Genetics 91, 67–94 (1979).

29. Hu, C. et al. The Inner Nuclear Membrane Protein Bqt4 in Fission Yeast Contains a DNA-Binding Domain Essential for Telomere Association with the Nuclear Envelope. Structure 27, 335–343.e3 (2019).

30. Kim, Y., Kostow, N. & Dernburg, A. F. The Chromosome Axis Mediates Feedback Control of CHK-2 to Ensure Crossover Formation in C. elegans. Dev Cell 35, 247–261 (2015).

31. Kleckner, N. Chiasma formation: chromatin/axis interplay and the role(s) of the synaptonemal complex. Chromosoma 115, 175–194 (2006).

32. Labella, S., Woglar, A., Jantsch, V. & Zetka, M. Polo kinases establish links between meiotic chromosomes and cytoskeletal forces essential for homolog pairing. Dev Cell 21, 948–958 (2011).

33. Link, J. & Jantsch, V. Meiotic chromosomes in motion: a perspective from Mus musculus and Caenorhabditis elegans. Chromosoma 128, 317–330 (2019).

34. Link, J. et al. Transient and Partial Nuclear Lamina Disruption Promotes Chromosome Movement in Early Meiotic Prophase. Dev Cell 45, 212–225.e7 (2018).

35. Liu, C. et al. A cooperative network at the nuclear envelope counteracts LINC-mediated forces during oogenesis in C. elegans. broRxiv (2021)

36. MacQueen, A. J. et al. Chromosome sites play dual roles to establish homologous synapsis during meiosis in C. elegans. Cell 123, 1037–1050 (2005).

37. McKim, K. S., Howell, A. M. & Rose, A. M. The effects of translocations on recombination frequency in Caenorhabditis elegans. Genetics 120, 987–1001 (1988).

38. Mikolcevic, P. et al. Essential role of the Cdk2 activator RingoA in meiotic telomere tethering to the nuclear envelope. Nature Communications 7, 11084 (2016).

39. Page, S. L. & Hawley, R. S. The genetics and molecular biology of the synaptonemal complex. Annu Rev Cell Dev Biol 20, 525–558 (2004).

40. Palmer, N. et al. A novel function for CDK2 activity at meiotic crossover sites. PLoS Biol 18 (2020).

41. Penkner, A. M. et al. Meiotic chromosome homology search involves modifications of the nuclear envelope protein Matefin/ SUN-1. Cell 139, 920–933 (2009).

42. Penkner, A. et al. The nuclear envelope protein Matefin/SUN-1 is required for homologous pairing in C. elegans meiosis. Dev Cell 12, 873–885 (2007).

43. Phillips, C. M. & Dernburg, A. F. A family of zinc-finger proteins is required for chromosome-specific pairing and synapsis during meiosis in C. elegans. Dev Cell 11, 817–829 (2006).

44. Phillips, C. M. et al. Identification of chromosome sequence motifs that mediate meiotic pairing and synapsis in C. elegans. Nat Cell Biol 11, 934–942 (2009).

45. Phillips, C. M. et al. HIM-8 binds to the X chromosome pairing center and mediates chromosome-specific meiotic synapsis. Cell 123, 1051–1063 (2005).

46. Rillo-Bohn, R. et al. Analysis of meiosis in Pristionchus pacificus reveals plasticity in homolog pairing and synapsis in the nematode lineage. Elife 10 (2021).

47. Rog, O. & Dernburg, A. F. Chromosome pairing and synapsis during Caenorhabditis elegans meiosis. Curr Opin Cell Biol 25, 349–356 (2013).

48. Rosenbluth, R. E. & Baillie, D. L. The genetic analysis of a reciprocal translocation, eT1(III; V), in Caenorhabditis elegans. Genetics 99, 415–428 (1981).

49. Sato, A. et al. Cytoskeletal forces span the nuclear envelope to coordinate meiotic chromosome pairing and synapsis. Cell 139, 907–919 (2009).

50. Scherthan, H. A bouquet makes ends meet. Nat Rev Mol Cell Biol 2, 621–627 (2001).

51. Scherthan, H. et al. Centromere and telomere movements during early meiotic prophase of mouse and man are associated with the onset of chromosome pairing. J Cell Biol 134, 1109–1125 (1996).

52. Shibuya, H. et al. MAJIN Links Telomeric DNA to the Nuclear Membrane by Exchanging Telomere Cap. Cell 173, 1058 (2018).

53. Spencer, W. C. et al. A spatial and temporal map of C. elegans gene expression. Genome Res 21, 325–341 (2011).

54. Swan, K. A. et al. High-throughput gene mapping in Caenorhabditis elegans. Genome Res 12, 1100–1105 (2002).

55. Tu, Z. et al. Speedy A–Cdk2 binding mediates initial telomere–nuclear envelope attachment during meiotic prophase I independent of Cdk2 activation. Proc Natl Acad Sci U S A 114, 592–597 (2017).

56. Viera, A. et al. CDK2 is required for proper homologous pairing, recombination and sex-body formation during male mouse meiosis. J Cell Sci 122, 2149–2159 (2009).

57. Villeneuve, A. M. A cis-acting locus that promotes crossing over between X chromosomes in Caenorhabditis elegans. Genetics 136, 887–902 (1994).

58. Wang, G. et al. Tethering of Telomeres to the Nuclear Envelope Is Mediated by SUN1-MAJIN and Possibly Promoted by SPDYA-CDK2 During Meiosis. Front Cell Dev Biol 8 (2020).

59. Wang, Y. et al. The meiotic TERB1-TERB2-MAJIN complex tethers telomeres to the nuclear envelope. Nat Commun 10, 564 (2019).

60. Wicks, S. R., Yeh, R. T., Gish, W. R., Waterston, R. H. & Plasterk, R. H. Rapid gene mapping in Caenorhabditis elegans using a high density polymorphism map. Nat Genet 28, 160–164 (2001).

61. Woglar, A. et al. Matefin/SUN-1 phosphorylation is part of a surveillance mechanism to coordinate chromosome synapsis and recombination with meiotic progression and chromosome movement. PLoS Genet 9, e1003335 (2013).

62. Woglar, A. & Villeneuve, A. M. Dynamic Architecture of DNA Repair Complexes and the Synaptonemal Complex at Sites of Meiotic Recombination. Cell 173, 1678–1691.e16 (2018).

63. Wynne, D. J., Rog, O., Carlton, P. M. & Dernburg, A. F. Dyneindependent processive chromosome motions promote homologous pairing in C. elegans meiosis. J Cell Biol 196, 47–64 (2012).

64. Zhang, L., Ward, J. D., Cheng, Z. & Dernburg, A. F. The auxininducible degradation (AID) system enables versatile conditional protein depletion in C. elegans. Development 142, 4374–4384 (2015).

65. Zhang, L. et al. Meiotic cell cycle progression requires adaptation to a constitutive DNA damage signal. broRxiv (2021)

66. Zickler, D. & Kleckner, N. Meiotic chromosomes: integrating structure and function. Annu Rev Genet 33, 603–754 (1999).

67. Zickler, D. & Kleckner, N. The leptotene-zygotene transition of meiosis. Annu Rev Genet 32, 619–697 (1998).

68. Zickler, D. & Kleckner, N. Recombination, Pairing, and Synapsis of Homologs during Meiosis. Cold Spring Harb Perspect Biol 7 (2015).

